# Void spot assay visualization optimization for use of Void Whizzard in rats (*Rattus norvegicus*)

**DOI:** 10.1101/2025.01.14.633030

**Authors:** Hannah Ruetten, Jadyn Bothe, Shannon Lankford, Gopal Badlani, James Koudy Williams

**Author notes:** **Corresponding Author:** Hannah Ruetten, Wake Forest Institute for Regenerative Medicine, 391 Technology Way, Winston-Salem, NC, 27157, USA. H. Ruetten and J. Bothe contributed equally to this work.

## Abstract

Void spot assay (VSA) noninvasively evaluates urination. VSA is often not performed in rats due to difficulty analyzing larger papers compared with mouse. This study optimizes VSA for rats by comparing post-assay visualization techniques: bright field light (BF), ultraviolet light (UV), and ninhydrin spray (N). Male rats were placed in cages lined by filter paper for 4 hours. Water and food was provided ad lib. After the assay all papers were dried overnight. BF images were photographed (digital camera). UV images were captured after cutting papers in half using a Darkroom ultraviolet imaging cabinet. Papers were sprayed with ninhydrin and photographed (digital camera). All images were converted to black and white for analysis with Void Whizzard. UV vs. BF showed significant differences in area. All three groups had significant differences in overall spot count and in the smallest sized spots (0-0.1 cm^2^). UV vs. N and UV vs. BF showed significant differences in 0.1-0.25 cm^2^ spots, UV vs. N in 0.25-0.5 cm^2^, and N vs. BF in spots 0.5-1 cm^2^. Overall BF visualization proved difficult. N provided an ideal way to highlight urine and image with a digital camera. Human fingerprints from pre-assay handling of paper interfered with analysis of the smallest sized spots however there were no differences in detection of larger spots, spot distribution, or overall spot area. This study contributes to the development of a standardized VSA protocol for assessing bladder function in both mouse and rat models.

## Introduction

A number of rat models exist for urinary disorders including bladder outlet obstruction, incontinence, pelvic floor dysfunction, spinal cord injury, and diabetic urinary dysfunction (1-3). Rats are often preferred over mice when model creation or treatment interventions involve surgery due to their increased size. Many labs perform cystometry on rats or utilize metabolic cages to assess urinary function however these methods require expensive equipment and at minimum involve removing the rat from its home cage which could impact urinary behavior due to handling stress and scent marking. In the case of cystometry, an invasive surgical procedure to implant a catheter in the bladder is performed which impacts urinary function.

A common urinary function assessment performed in awake mice in their home cage without any specialized equipment or surgical manipulation is the void spot assay. In the void spot assay a mouse is placed in an enclosure lined with paper, they move freely while micturition is captured. The paper is then removed, imaged under UV light, and fed into an ImageJ plugin called “Void Whizzard” which traces the spots and gives you a readout of spot size, spot count, and spot distribution. The assay in mice has been extensively optimized, software has been developed for quick and easy assessment, and the assay has been shown to have good predictability not only in showing changes in urinary function over time but also in discriminating differing voiding phenotypes from one another (4-8).

Void spot assays are typically not performed in rats due to difficulty in analyzing papers from larger rat cages. However some have performed “urine-marking tests” in rats by hand drawing/tracing urine spots and performing manual measurements (9). In a study by Wegner et al. it was shown that laboratory-specific manual methods for void spot assay image analysis cause substantial variability in end point measurements (8). Therefore it would be ideal to optimize a way to image void spot assay papers from rats and use the images for analysis with Void Whizzard. This study aims to improve feasibility of VSA for rats by testing agents to alter urine color and comparing post-assay visualization techniques: bright field light, ultraviolet light, and ninhydrin spray.

## Materials and Methods

### Rats

All experiments were conducted under an approved protocol from the Wake Forest University Animal Care and Use Committee and in accordance with the National Institutes of Health *Guide for the Care and Use of Laboratory Animals*. Heterogenous stock rats (n=32) were acquired as overstock from a campus breeding colony. Rats were housed in Allentown square microisolator rat caging on an Allentown rack (Allentown, Allentown, NJ). Room lighting was maintained on a 12:12-h light-dark cycles, room temperature was typically 20.5 ± 5°C, and humidity was 30–70%. Rats were fed 5P76 - Prolab® IsoPro® RMH 3000, and food and water were available ad libitum. Cages contained Bed-o’cob® ¼” bedding. All papers were collected at the baseline of the study from unmanipulated rats.

### Color Altering Agents

Two rats were provided 200mg of phenazopyridine dissolved in drinking water for 4 hours and water consumption was measured. The dose was halved to 100mg and tested for the same duration in two other rats. Animal resources was contacted and several combinations of phenazopyridine and sweeteners (honey, dextrose, and glucose) were trialed and consumption measured.

Rats were tested for consumption of beets and raspberries. Four rats were provided with two dishes per cage (1/rat) containing 4-6 cubes (5-15mm at the largest dimension) of raw beetroot and 1 red raspberry at 2PM the day prior to the void spot assay test. The test was repeated providing the beets and raspberry at 5PM the day prior to void spot assay in two additional rats. Finally, all 32 rats were provided beets and a raspberry at 5PM the day prior to the void spot assay test.

### Void Spot Assay

We followed the recommended guidelines of reporting VSA data (5, 6, 8). VSA was performed in the vivarium on the rack where the rats were housed. Cytiva Whatman™ 3MM Chr Chromatography Paper (26 × 41 cm sheets, pk 100) (no. No.05-713-336, Fisher Scientific) were purchased and cut with a paper cutter to match the floor dimensions of the rats’ home cage (26 × 36 cm). The paper was placed in the bottom of the cage with a nylabone placed in the center to discourage chewing. Rats were placed in the cage (singly housed) with food and water ad libitum for 4 h starting from 9 to 10 AM Eastern time.

### Ultraviolet Light Paper Imaging

Filter papers were dried then cut in half and imaged with an Autochemi AC1 Darkroom ultraviolet imaging cabinet (UVP, Upland, CA) equipped with an Auto Chemi Zoom lens 2UV and an epi-illuminator. Image capture settings were adjusted using UVP VisonWorksLS image acquisition software. Images were captured using an Ethidium Bromide filter set (570–640 nm) and 365-nm epi-illumination (Figure 1A). Raw .tif files of each half of the paper were recombined using GIMP 2.10.36 (revision 1). Any chewed areas were manually filled using white color with the paint brush in JS Paint version 1.0.0+ (Figure 1B). Final images were cropped to the paper boundaries, and a macro utilized (run(“16-bit”); //run(“Brightness/Contrast…”); run(“Enhance Contrast”, “saturated=0.35”); run(“Apply LUT”)) to enhance contrast and save as 16-bit for analysis using Void Whizzard (Figure 1C).

**Figure 1.**
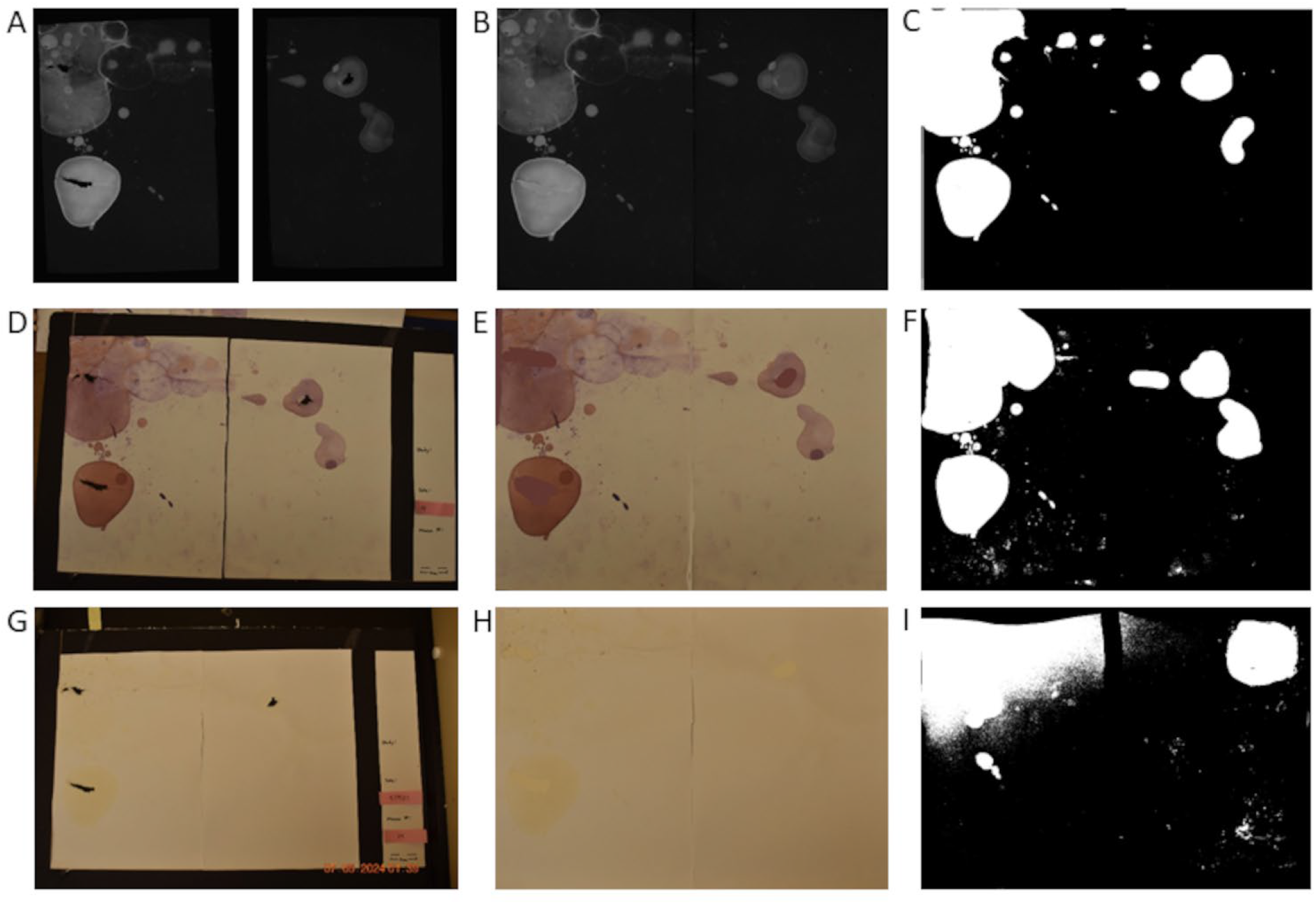
A Comparison of the Three Imaging Methods. (A) Ultraviolet light raw images from Autochemi AC1 Darkroom ultraviolet imaging cabinet, (B) raw images combined together, and (C) final black and white image for void spot assay analysis. (D) Picture of ninhydrin sprayed paper from digital camera, (E) image cropped and edited with chew spots filled in, and (F) final black and white image for void spot assay analysis. (G) Picture of void spot assay paper from digital camera, (H) image cropped and edited with chew spots filled in, and (I) final black and white image for void spot assay analysis.

### Bright field Paper Imaging

An imaging board was constructed using black foam board from the Dollar Tree. A large section of foam boarded was cut to a few inches larger than the VSA paper. Thin strips were then cut and layered on top at the left and bottom edge so there was a rigid L-shaped ledge to place the paper on. On the right side of the board a sheet of paper was placed with spaces to put the paper information to be recorded in the paper image. A scale bar was added so it was automatically included in the image. Papers were imaged with a Nikon digital camera (Figure 1D). Captured images were relabeled with the paper information and cropped to the paper margins. Any chewed areas were manually filled using light yellow color with the paint brush tool in JS Paint version 1.0.0+ (Figure 1E). Multiple techniques were trialed using ImageJ, including the threshold feature and the Colour Deconvolution2 tool, to separate the urine spots from the white paper background. Utilizing the Color Deconvolution2 plugin, ROI vectors were selected, with the first color set to blue for optimal contrast, the second “color matched” to a urine spot, and the third color “color matched” to the paper. For thresholding, the process involved navigation to Image > Adjust > Color Threshold. In the popup window, the “dark background” option was deselected, and the default thresholding method, white threshold color, and HSB color space were used. The following parameters were applied: Hue (pass deselect, 71-105), Saturation (select pass, 0-209), and Brightness (select pass, 0-97).Ultimately the thresholding method was determined to be the most effective and used in final analysis. Final images were cropped to the paper boundaries, and converted to 16bit .tiff files for analysis using Void Whizzard (Figure 1F).

### Ninhydrin Paper Imaging

Ninhydrin Aerosol Spray- 16 oz (A-2643, Thomas Scientific) was used to apply a light coat of spray to each urine paper. The papers were left to dry overnight. The same imaging board used for Bright field was used for Ninhydrin. Papers were imaged with a Nikon digital camera (Figure 1G). Captured images were relabeled with the paper information and cropped to the paper margins. Any chewed areas were manually filled using light purple color with the paint brush tool in JS Paint version 1.0.0+ (Figure 1H). Multiple techniques were trialed using ImageJ, including the threshold feature and Colour Deconvolution2 tool, to separate the urine spots from the background white paper. For the Color Deconvolution2 plugin, ROI vectors were selecting with the following parameters: the first color was set to yellow to achieve the greatest contrast against the purple ninhydrin spots, the second color was matched to a urine spot, and the third color was matched to the paper. For the thresholding technique, (Image > Adjust > Color Threshold) in the popup window, “dark background” option was deselected, and the default thresholding method, black and white threshold color, and lab color space were used. The following parameters were applied: L*(pass selected, 0-84), a* (pass selected, (0-255), and b* (pass selected, 0-255). The thresholding method was determined to be the most effective and used for final analysis. Final images were cropped to the paper boundaries, and converted to 16bit .tiff files for analysis using Void Whizzard (Figure 1I).

### Void Paper Analysis

Void Whizzard was downloaded from http://imagej.net/Void_Whizzard and run according to the user guide (8). Analyzed parameters included total spot count, total void area (cm2), percent area in the center of the paper, percent area in corners of the paper, and mass distribution of spots (0–0.1, 0.1– 0.25, 0.25–0.5, 0.5–1, 1–2, 2–3, 3–4, and 4+ cm2).

### Statistical Analysis

Statistical analyses were performed with Graph Pad Prism 8.0.2. Differences were considered significant at the P < 0.05 level. For group comparisons, a Kruskal-Wallis test was applied along with Dunn’s multiple comparisons test.

## Results and Discussion

### Rats Wouldn’t Drink Water Containing Phenazopyridine Even With Sweetener

Phenazopyridine (2,6-diamino-3-[phenyl-azo]pyridine) is used as an oral analgesic medication to treat patients with bladder inflammation. A universal side effect from taking this medication is that it turns the urine orange. Urologists in addition to using the drug for treating bladder inflammation have also leveraged this drug to color the urine orange, specifically for the purposes of cystoscopic identification of the ureteral orifices (10). In this study we attempted to color rat urine orange in order to allow for better bright field visualization of urine spots.

Rats were tested for consumption of phenazopyridine in drinking water. Two rats were provided 200mg of phenazopyridine dissolved in drinking water for 4 hours and water consumption was measured. Neither rat consumed any water with phenazopyridine. They both immediately drank regular water upon returning it to the cage. The dose was halved to 100mg and tested for the same duration in two other rats with the same results. Animal resources were contacted and several combinations of phenazopyridine and sweeteners (honey, dextrose, and glucose) were trialed with no success in rats consuming the water.

In prior studies of phenazopyridine pharmacokinetics in rats, researchers dosed the medication via oral gavage (11). However this would induce handling stress immediately prior to the void spot assay so was not pursued. Other researchers have dosed medications mixed into peanut butter and, anecdotally from a lab animal veterinary resident, rats seem to take medications well mixed into cream of chicken soup. Ultimately these routes were also not pursued due to need for handling and individual rat dosing.

If the phenazopyridine dosing had worked there would also likely have been drug related effects that would have masked some urinary changes. When dosed intravenously in female Sprague-Dawley rats phenazopyridine increases bladder compliance at 0.3, 1, and 3 mg/kg dose levels (12).

### Rats Would Consume Beets and Raspberries but the Resulting Urine Color Alteration Was Not Predictable and Durable for the Duration of the Void Spot Assay

Beeturia occurs in 13.8% of the human population and is characterized by a pink to dark red discoloration of the urine (13). In rats dosed with fermented beet juice betacyanin derivatives are noted in the urine as soon as 15min after administration and persist for at least two hours (14). In rats dosed with beetroot powder about 3% of the original betanin was recovered in the urine over the following 24 hours (15). Anecdotally rats that receive a lot of beets or red berries in their diet can develop red discoloration of the urine that owners mistake for hematuria. However, beet over consumption can lead to kidney stones (16) so beet intake must be kept within acceptable ranges set by animal resources. Based on this we hypothesized that providing our rats with a moderate sized treat of beets and red raspberries the night before void spot assay may discolor the urine pink to dark red and aid in void spot visualization.

Rats were tested for consumption of beets and raspberries. Four rats were provided two dishes per cage (1/rat) containing 4-6 cubes (5-15mm at the largest dimension) of raw beetroot and 1 red raspberry at 2PM the day prior to the void spot assay test. Rats readily consumed the beets and raspberries. Void spot assay was performed the following morning at 9 AM and the urine took on a slight pink hue in a few voids for the rats but returned to normal color by the end of the assay. The test was repeated providing the beets and raspberry at 5PM the day prior to void spot assay in two additional rats with similar results. All 32 rats were provided beets and a raspberry at 5PM the day prior to the void spot assay test. Some voids had a slight pink hue but quickly faded to normal urine color before the papers could be analyzed.

The fading of the pink urine hue during paper drying is likely related to it becoming more alkaline. This also occurs with human urine samples which are darker pink when they are acidic but the discoloration goes away as they become alkaline (13). Overall the rats enjoyed their treat but it did not provide the desired result of aiding in void bright field visualization.

### Papers Analyzed With the Ninhydrin Method Have a Similar Total Void Area, Spot Distribution, and Number of Larger Spots to Ultraviolet Light Method

Since we were unable to color the urine in the rats, we next tested a method to color the urine after it was deposited on the paper. In a study in 2008, Arakawa et al. used Ninhydrin spray to detect urine from mouse scent marking assays (17). Ninhydrin is one of the most popular and widely used reagents for detecting proteins. It is used in a range of protein detection reactions, but one of its most commonly known uses is detection of finger prints. Because of its strong color changing power in the presence of protein it readily changes urine from a light yellow to a vibrant purple color. Here we compare visualization with ninhydrin to both ultraviolet light urine imaging and bright field imaging without any color altering aid.

The first set of parameters we compared were total void area, total void count, percent void area in center, and percent void area in corners. The ninhydrin method had a similar total void area (p>0.9999) (Figure 2A), percent void area in center(p>0.9999) (Figure 2C), and percent void area in corner (p>0.9999) (Figure 2D) compared to ultraviolet method. Bright field method had a higher total void area (p=0.0011, p<0.0001) (Figure 2A), similar percent void area in center (p>0.9999, p>0.9999) (Figure 2C), and decreased void area in corners (p=0.0263, p=0.0019) (Figure 2D) compared to ultraviolet and ninhydrin method. Total spot count was different for each method; it was slightly higher for ninhydrin (p<0.0001), and much higher for bright field (p<0.0001) compared to ultraviolet method (Figure 2B).

**Figure 2.**
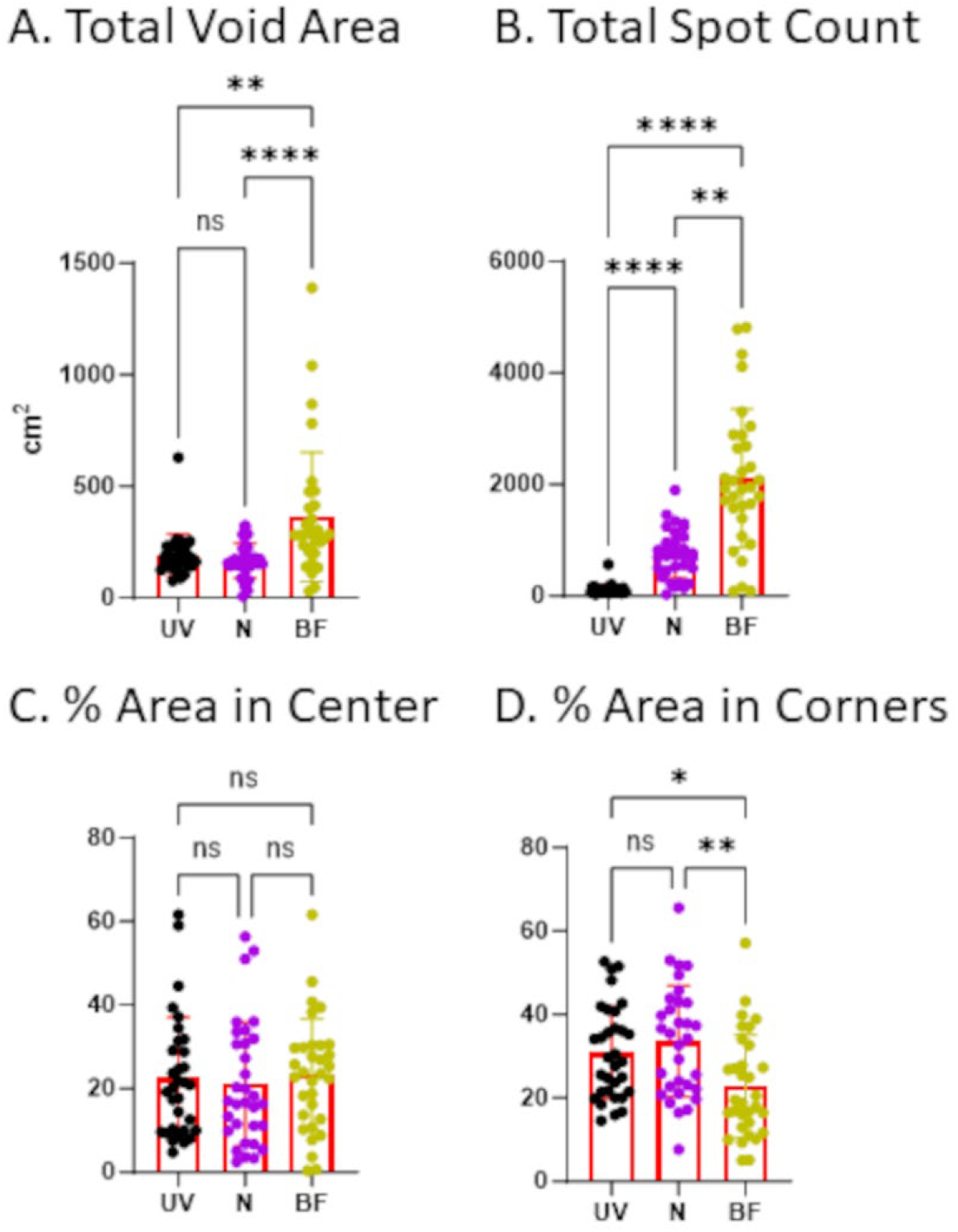
Papers Analyzed With the Ninhydrin Method Have a Similar Total Void Area and Spot Distribution to Ultraviolet Light Method. Male rats (n=32) were placed in cages lined by filter paper for 4 hours. Water and food was provided ad lib. After the assay all papers were dried overnight. Bright field (BF) images were photographed using a digital camera. Ultraviolet light (UV) images were captured after cutting papers in half using a Darkroom ultraviolet imaging cabinet. Papers were sprayed with ninhydrin (N) and photographed (digital camera). All images were converted to black and white for analysis with Void Whizzard. Statistical analyses were performed with Graph Pad Prism 8.0.2. Differences were considered significant at the P < 0.05 level. For group comparisons, a Kruskal-Wallis test was applied along with Dunn’s multiple comparisons test. (ns) p-value >0.05, (*) p-value <0.05 but >0.01, (**) p-value <0.01, (***) p-value <0.001, (****) p-value <0.0001.

The next set of parameters we compared were the small spots: 0-0.1cm^2^ spots, 0.1-0.25cm^2^ spots, 0.25-0.5cm^2^ spots, 0.5-1cm^2^ spots, and 1-2cm^2^ spots. The number of 0-0.1cm^2^ spots were different for each method; it was slightly higher for ninhydrin (p<0.0001), and much higher for bright field (p<0.0001) compared to ultraviolet method (Figure 3A). Both ninhydrin (p=0.0002) and bright field (p=0.0042) had more 0.1-0.25cm^2^ spots than ultraviolet method (Figure 3B). For the ninhydrin method we believe these increases in tiny spots were due to small rat footprints being detected as small urine spots. For bright field method we believe these increases in tiny spots were due to shadow artifact and dust on the paper being detected as small urine spots. The number of 0.25-0.5cm^2^ spots were slightly higher for ninhydrin than bright field method (p=0.0013) and ultraviolet method (p=0.0072); bright field and ultraviolet were similar (p>0.9999) (Figure 3C). The number of 0.5-1cm^2^ spots were similar between ultraviolet method and ninhydrin method (p=0.0737); bright field method was similar to ultraviolet method (p=0.2825) but lower than ninhydrin (p=0.0003) (Figure 3D). The number of 1-2cm^2^ spots were slightly higher for ninhydrin than bright field method (p=0.0026) and ultraviolet method (p=0.0164); bright field and ultraviolet were similar (p>0.9999) (Figure 3E). We believe these increases in 0.25-0.5cm^2^ and 1-2cm spots in ninhydrin compared to ultraviolet were due to partial and whole human fingerprints being detected as urine spots.

**Figure 3.**
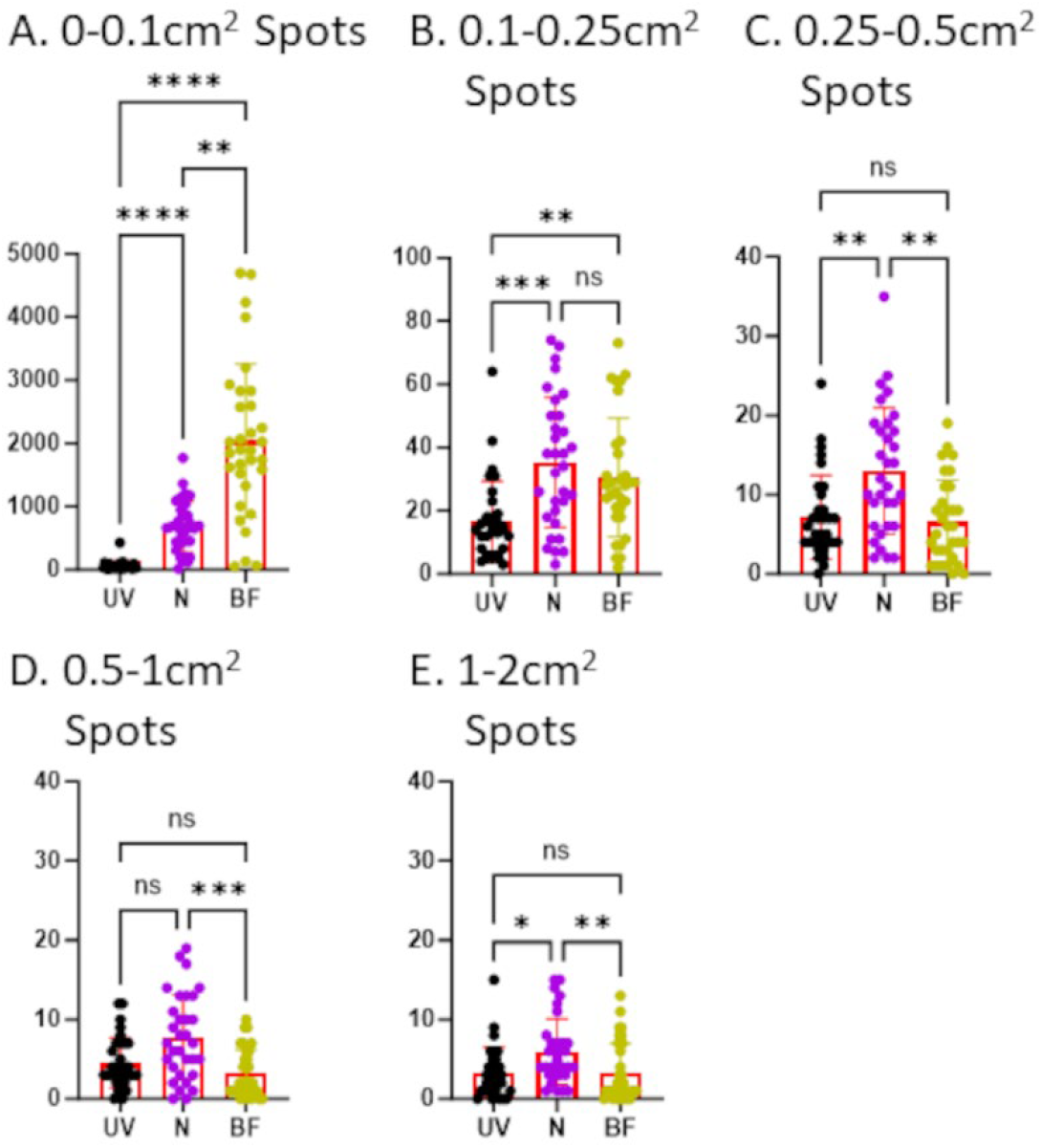
Papers Analyzed With the Ninhydrin Method Have More Small Spots Compared to Ultraviolet Light Method. Male rats (n=32) were placed in cages lined by filter paper for 4 hours. Water and food was provided ad lib. After the assay all papers were dried overnight. Bright field (BF) images were photographed using a digital camera. Ultraviolet light (UV) images were captured after cutting papers in half using a Darkroom ultraviolet imaging cabinet. Papers were sprayed with ninhydrin (N) and photographed (digital camera). All images were converted to black and white for analysis with Void Whizzard. Statistical analyses were performed with Graph Pad Prism 8.0.2. Differences were considered significant at the P < 0.05 level. For group comparisons, a Kruskal-Wallis test was applied along with Dunn’s multiple comparisons test. (ns) p-value >0.05, (*) p-value <0.05 but >0.01, (**) p-value <0.01, (***) p-value <0.001, (****) p-value <0.0001.

The final set of parameters we compared were the large spots: 2-3cm^2^ spots, 3-4cm^2^ spots, and 4+cm^2^. For all three spot sizes the number of spots detected were similar among all three methods (p=0.1329->0.9999) (Figure 4).

**Figure 4.**
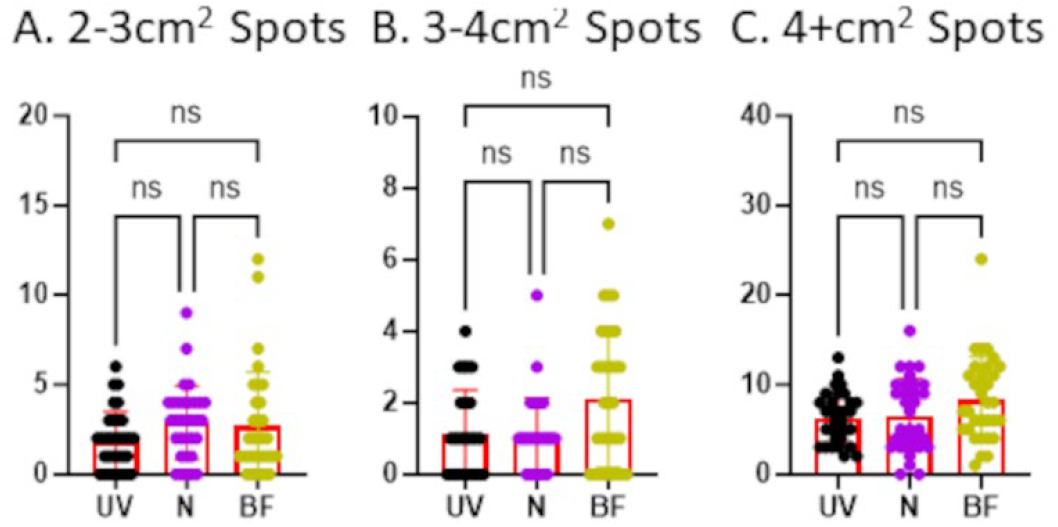
All three methods have similar numbers of large spots. Male rats (n=32) were placed in cages lined by filter paper for 4 hours. Water and food was provided ad lib. After the assay all papers were dried overnight. Bright field (BF) images were photographed using a digital camera. Ultraviolet light (UV) images were captured after cutting papers in half using a Darkroom ultraviolet imaging cabinet. Papers were sprayed with ninhydrin (N) and photographed (digital camera). All images were converted to black and white for analysis with Void Whizzard. Statistical analyses were performed with Graph Pad Prism 8.0.2. Differences were considered significant at the P < 0.05 level. For group comparisons, a Kruskal-Wallis test was applied along with Dunn’s multiple comparisons test. (ns) p-value >0.05, (*) p-value <0.05 but >0.01, (**) p-value <0.01, (***) p-value <0.001, (****) p-value <0.0001.

## Conclusions

When researchers have access to the more specialized equipment, the UV method remains the superior method for imaging void spot assay papers. However, due to potential exposure of humans to high levels of UV when imaging these large papers, expense of equipment, and added time to cut apart, image, and rebuild full images of the papers, N provided an ideal alternative. N spray is relatively inexpensive and allows you to image the papers with any camera. Human fingerprints from pre-assay handling of paper as well as rat footprints interfered with analysis of the smallest sized spots however human fingerprint interference is easily minimized by use of gloves when handling the paper pre- and post-assay. Additionally with the N method there were no differences in detection of the larger spots, spot distribution, or overall spot area. This study contributes to the development of a standardized VSA protocol for assessing bladder function in both mouse and rat models.

**Table 1.** Void Spot Assay Results Summary.

| Parameter | Ultraviolet (UV)<br>n=32<br>(mean ± SD) | Ninhydrin (N)<br>n=32<br>(mean ± SD) | Brightfield (BF)<br>n=32<br>(mean ± SD) | UV vs N<br>(p-value) | UV vs BF<br>(p-value) | N vs BF<br>(p-value) |
| --- | --- | --- | --- | --- | --- | --- |
| **Total Void Area** | 192.1 ± 93.99 | 166.2 ± 78.49 | 363.5 ± 288.7 | $>0.9999$ | **0.0011** | **<0.0001** |
| **Total Spot Count** | 108.7 ± 92.56 | 751.0 ± 434.5 | 2103 ± 1242 | **<0.0001** | **<0.0001** | **0.0030** |
| **% Center** | 22.57 ± 14.44 | 21.08 ± 14.76 | 23.32 ± 13.41 | $>0.9999$ | $>0.9999$ | $>0.9999$ |
| **% Corners** | 30.85 ± 11.11 | 33.57 ± 13.28 | 22.74 ± 12.30 | $>0.9999$ | **0.0263** | **0.0019** |
| **0-0.1cm<sup>2</sup> spots** | 67.81 ± 72.52 | 678.4 ± 402.7 | 2046 ± 1215 | **<0.0001** | **<0.0001** | **0.0025** |
| **0.1-0.25cm<sup>2</sup> spots** | 16.56 ± 12.77 | 35.25 ± 20.50 | 30.47 ± 18.74 | **0.0002** | **0.0042** | $>0.9999$ |
| **0.25-0.5cm<sup>2</sup> spots** | 7.156 ± 5.292 | 12.97 ± 7.966 | 6.563 ± 5.242 | **0.0072** | $>0.9999$ | **0.0013** |
| **0.5-1cm<sup>2</sup> spots** | 4.531 ± 3.152 | 7.781 ± 5.326 | 3.219 ± 3.003 | 0.0737 | 0.2825 | **0.0003** |
| **1-2cm<sup>2</sup> spots** | 3.313 ± 3.167 | 5.969 ± 4.115 | 3.250 ± 3.742 | **0.0164** | $>0.9999$ | **0.0026** |
| **2-3cm<sup>2</sup> spots** | 1.875 ± 1.641 | 2.875 ± 2.044 | 2.719 ± 2.986 | 0.1329 | $>0.9999$ | 0.7185 |
| **3-4cm<sup>2</sup> spots** | 1.156 ± 1.194 | 1.094 ± 1.058 | 2.125 ± 1.930 | $>0.9999$ | 0.1436 | 0.1468 |
| **4+cm<sup>2</sup> spots** | 6.313 ± 2.788 | 6.594 ± 4.047 | 8.375 ± 4.730 | $>0.9999$ | 0.2141 | 0.3563 |

## Acknowledgements

Bothe supported by NIH R25 DK121572; Ruetten supported by NIH T32 OD010957; Research supported by a grant from The Veterans Association Merit Award. The funding sources had no influence over study design, execution, interpretations, or development of the abstract.

## Author Contributions

Conceived and designed research (H.R.), performed experiments (H.R., S.S.L., J.B.,), analyzed data (J.B. and H.R.), interpreted results of experiments (J.B. and H.R.), prepared figures (J.B. and H.R.), drafted manuscript (H.R.), edited and revised manuscript (H.R., J.B., S.S.L., G.B., and J.K.W.), approved final version of manuscript (H.R., J.B., S.S.L., G.B., and J.K.W.).

